## Supplementary material for "Single-cell ATAC-seq Signal Extraction and Enhancement with SCATE": Additional file 1 - Table S1.pdf

**Table S1**. A comparison between SCATE and other existing methods

| Method | Combine CREs | Combine cells | Adaptively tune resolution | Use public bulk data to model baseline | Use Binary or Count data | Primary goal | Reference |
|---|---|---|---|---|---|---|---|
| SCATE | √ | √ | √ | √ | C | Reconstruct activities of each individual CRE | This paper |
| chromVAR | √ | | | | C | Cluster cells, identify TF motifs associated with differential accessibility and variability | [12] |
| SCRAT | √ | | | | C | Cluster cells, identify CRE pathways associated with differential accessibility | [13] |
| BROCKMAN | √ | | | | B | Summarize data by k-mers and perform principal component analysis on k-mer features to identify co-varying TFs, cluster cells | [14] |
| Dr.seq2 | | √ | | | C | Cluster cells, identify peaks (MACS) in each cell subpopulation | [17] |
| Cicero | | √ | | | B | Identify correlated pairs of CREs | [18] |
| Scasat | | | | | B | Cluster cells, identify peaks (MACS), differential accessibility analysis | [20] |
| Destin | | | | | B | Cluster cells | [21] |
| scABC | | | | | C | Cluster cells | [22] |
| PRISM | | | | | B | Quantify cell-to-cell variation to identify hyper- or hypo-variable genomic features | [23] |
| cisTopic | | | | | B | Represent data using low-dimensional topic-cell and region-topic representation, cluster cells and CREs accordingly | [24] |
