## Supplementary material for "Single-cell ATAC-seq Signal Extraction and Enhancement with SCATE": Additional file 2 - Figure S1-S3.pdf

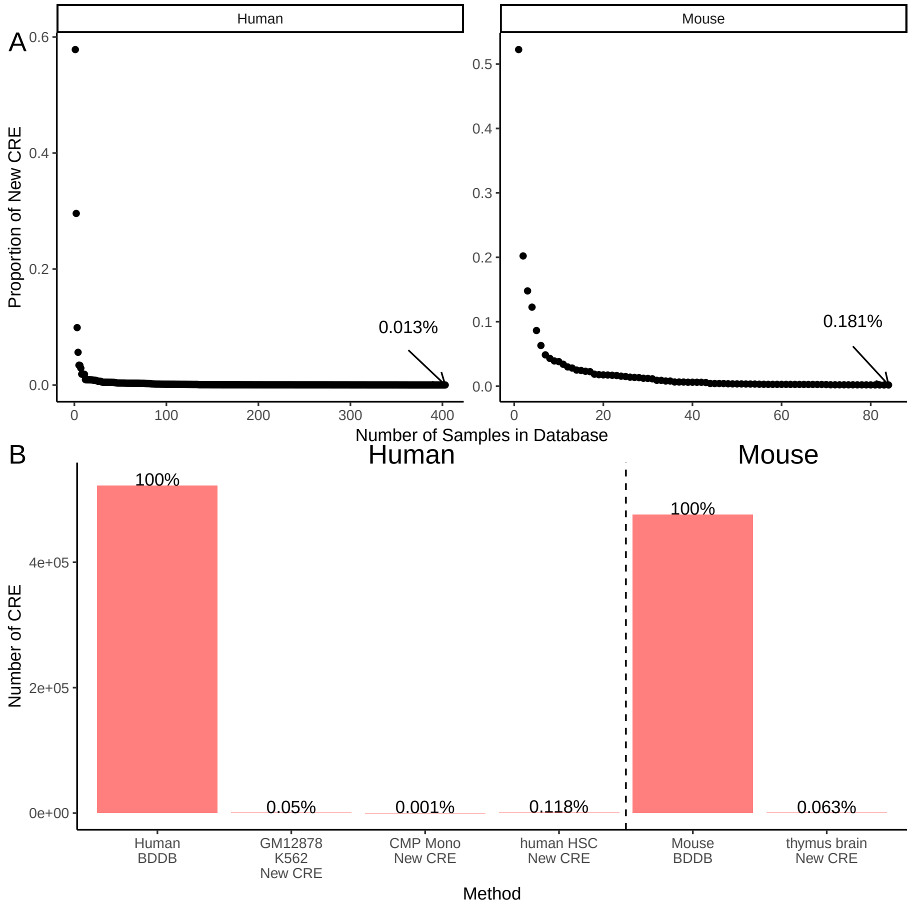


**Figure S1**: Saturation analysis of BDDB CRE lists. (A): As one increases the number of DNase-seq samples in the BDDB database, the proportion of new CREs contributed by adding a new sample gradually decreases. (B): The scATAC-seq datasets analyzed in this study would only add 0.0013%-0.118% new CREs to the precompiled CRE list in BDDB.

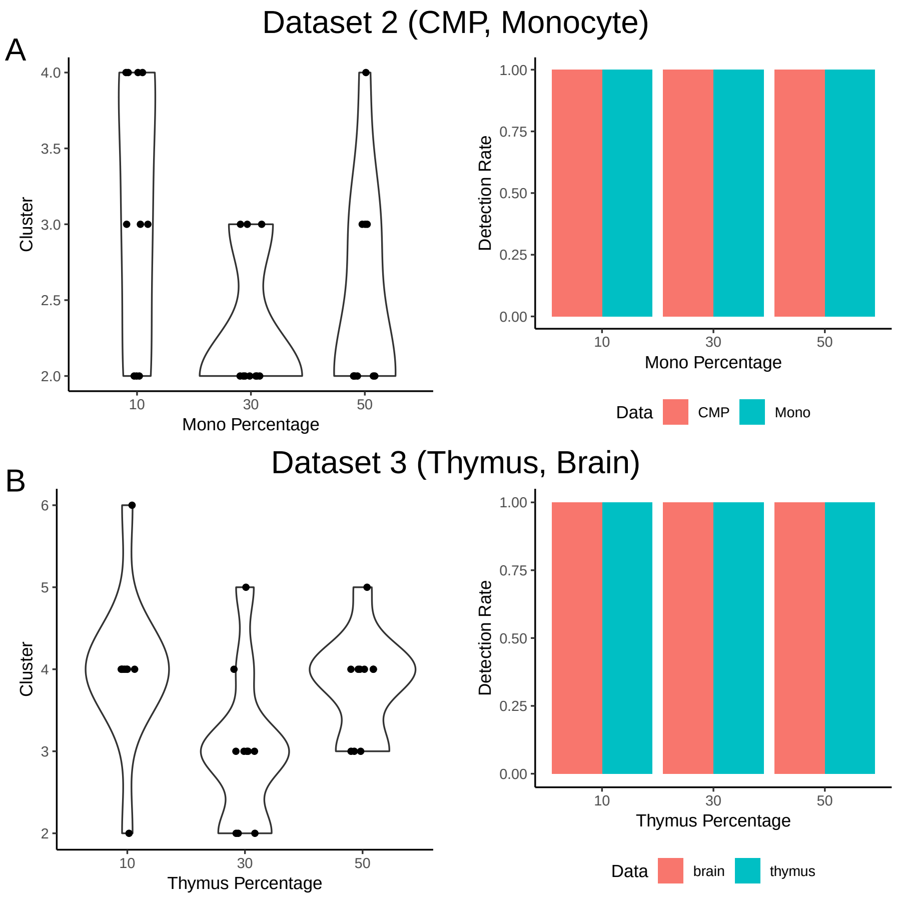


**Figure S2**: Analyses of a heterogeneous cell population created using (A) Dataset 2 and (B) Dataset 3. In each dataset, the left plot shows distribution of cell cluster numbers obtained by SCATE for synthetic samples with different cell mixing proportions. For each mixing proportion, 10 synthetic samples were created and analyzed. The right plot shows the frequency that each cell type is detected in the 10 synthetic samples at each cell mixing proportion.

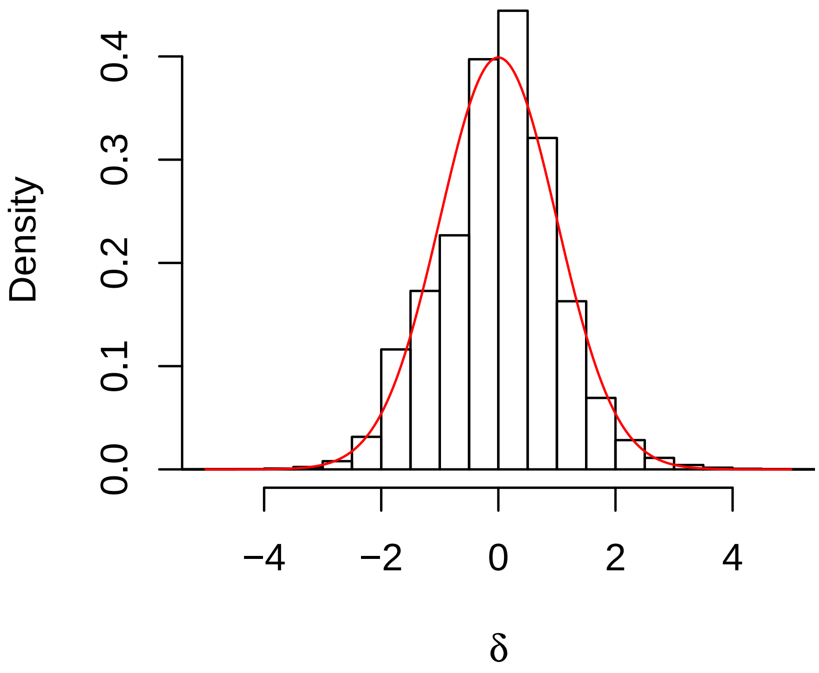


**Figure S3**: The empirical distribution (histogram) of the log-normalized read counts in human BDDB after standardization (i.e., subtract the mean and divide by SD of each CRE) can be fitted well with a normal distribution (red curve).
