## Supplementary material for "Single-cell ATAC-seq Signal Extraction and Enhancement with SCATE": Additional file 6 - Supplementary Note.pdf

### Additional file 6 – derivation of moment estimators for $m_i$ and $s_i$

Assume the read count $y_{i,j}$ for the $i$-th CRE and the $j$-th bulk DNase-seq sample is generated by a Poisson distribution:

$$\begin{aligned} y_{i,j} &\sim Poisson(L_j \mu_{i,j}) \\ \log(\mu_{i,j}) &= m_i + s_i \delta_{i,j} \end{aligned}$$

Further assume $\delta_{i,j} \sim N(0,1)$. According to the law of total expectation and total variance for conditional probability:

$$\begin{aligned} \mathrm{E}\left(\frac{y_{i,j}}{L_j}\right) &= \frac{1}{L_j}\mathrm{E}[\mathrm{E}(y_{i,j}|\delta_{i,j})] \\ &= \mathrm{E}[\mu_{i,j}] \\ &= \mathrm{E}[e^{m_i + s_i\delta_{i,j}}] \\ &= e^{m_i}\mathrm{E}[e^{s_i\delta_{i,j}}] \\ &= e^{m_i + \frac{1}{2}s_i^2} \end{aligned}$$

$$\begin{aligned} \mathrm{E}\left(\frac{y_{i,j}}{L_j}\right)^2 &= \frac{1}{L_j^2}\mathrm{E}[\mathrm{E}(y_{i,j}^2|\delta_{i,j})] \\ &= \frac{1}{L_j^2}\mathrm{E}[Var(y_{i,j}|\delta_{i,j}) + \mathrm{E}(y_{i,j}|\delta_{i,j})^2] \\ &= \frac{1}{L_j^2}\mathrm{E}[L_j\mu_{i,j} + L_j^2\mu_{i,j}^2] \\ &= \frac{1}{L_j}\mathrm{E}[\mu_{i,j}] + \mathrm{E}[\mu_{i,j}^2] \\ &= \frac{1}{L_j}e^{m_i + \frac{1}{2}s_i^2} + \mathrm{E}[e^{2m_i + 2s_i\delta_{i,j}}] \\ &= \frac{1}{L_j}e^{m_i + \frac{1}{2}s_i^2} + e^{2m_i + 2s_i^2} \\ &= \frac{1}{L_j}e^{m_i + \frac{1}{2}s_i^2} + \left[e^{m_i + \frac{1}{2}s_i^2}\right]^2 e^{s_i^2} \\ &= \frac{1}{L_j}\mathrm{E}\left(\frac{y_{i,j}}{L_j}\right) + \left[\mathrm{E}\left(\frac{y_{i,j}}{L_j}\right)\right]^2 e^{s_i^2} \end{aligned}$$

Thus we have:

$$e^{s_i^2} = \frac{\mathrm{E}\left(\frac{y_{i,j}}{L_j}\right)^2 - \frac{1}{L_j}\mathrm{E}\left(\frac{y_{i,j}}{L_j}\right)}{\left[\mathrm{E}\left(\frac{y_{i,j}}{L_j}\right)\right]^2}$$

and

$$s_i^2 = \log\left(\frac{\mathrm{E}\left(\frac{y_{i,j}}{L_j}\right)^2 - \frac{1}{L_j}\mathrm{E}\left(\frac{y_{i,j}}{L_j}\right)}{\left[\mathrm{E}\left(\frac{y_{i,j}}{L_j}\right)\right]^2}\right)$$

After replacing $\mathrm{E}\left(\frac{y_{i,j}}{L_j}\right)^2$ and $\mathrm{E}\left(\frac{y_{i,j}}{L_j}\right)$ by their sample estimates $\sum_j \left(\frac{y_{i,j}}{L_j}\right)^2/J$ and $\sum_j \left(\frac{y_{i,j}}{L_j}\right)/J$, respectively, we have:

$$\tilde{s}_i = \sqrt{\log\left(\frac{\sum_j (y_{i,j}/L_j)^2/J - \sum_j (y_{i,j}/L_j^2)/J}{(\sum_j (y_{i,j}/L_j)/J)^2}\right)}$$

Since

$$m_i = \log \mathrm{E}\left(\frac{y_{i,j}}{L_j}\right) - \frac{1}{2}s_i^2$$

We have

$$\tilde{m}_i = \log\left(\frac{\sum_j (y_{i,j}/L_j)}{J}\right) - \tilde{s}_i^2/2$$
